# Microbiome-behavior coupling shapes infant adaptation to early maternal unpredictability

**DOI:** 10.64898/2025.12.03.692149

**Authors:** Dima Amso, Guilherme Fahur Bottino, Tess Forest, Kevin S. Bonham, Michal R. Zieff, Fadheela Patel, Marlie Miles, Donna Herr, Khula South Africa Data Collection Team, Claudia Espinoza-Heredia, Celia L. D’Amato, Jinge Ren, Melissa Gladstone, Emmie Mbale, Daniel C. Alexander, Derek K Jones, Steve C. R. Williams, William P. Fifer, Laurel J. Gabard-Durnam, Vanja Klepac-Ceraj, Kirsten A. Donald

## Abstract

We asked whether infant microbiome composition, together with infants’ behavioral responses to early maternal unpredictability, can offer novel mechanistic insights into behavioral phenotypic variation in human development. Maternal unpredictability was computed as the entropy rate of transitions among maternal sensory signals during naturalistic mother-infant interaction (*n*=256 dyads; 2-6 months). Infant visual orienting behavior (VOB) was indexed by infants’ simultaneous gaze shifts to and from the mother. Higher maternal unpredictability predicted more frequent VOB shifts, indicating a mature-for-age profile, and longitudinally forecast poorer inhibitory control at 19-28 months. A subgroup of infants with high maternal unpredictability and high VOB had inhibitory control outcomes closer to those of infants who experienced low maternal unpredictability, suggesting VOB strategy might buffer against the effects of high maternal unpredictability. In participants with metagenomic data (*n*=93), VOB was associated with taxonomic and functional gut microbial profiles along a *Bifidobacterium breve* to *Bifidobacterium longum* axis. Neuroactive gene-set enrichment analysis linked VOB to increase in tryptophan synthesis and glutamate synthesis genes. In contrast, maternal unpredictability showed depletions of GABA and tryptophan synthesis in the unique-effects model and, when VOB and the maternal unpredictability by VOB term were included, enrichment of acetate synthesis and quinolinic acid degradation. Notably, the maternal unpredictability by VOB interactions for tryptophan and glutamate were negative, indicating that the VOB-pathway coupling attenuates as unpredictability increases. Together, these findings support a framework in which maternal unpredictability is the environmental challenge, infant VOB expresses the behavioral strategy, and microbial metabolism modulates whether that strategy is biochemically feasible.

**Teaser:** Infant gut microbiome is linked to infant behavioral responses to maternal unpredictability, which in turn impacts how they learn to control their actions.

## Introduction

Experiences early in human postnatal life have an outsized impact on later cognitive and mental health. However, the influence of early life experiences on later outcomes is almost always probabilistic rather than deterministic (*1*). Phenotypic plasticity, the capacity of a single genotype to produce different phenotypes, provides a framework for understanding such variation in human cognition and development (*2*). The mechanisms underlying phenotypic plasticity are certainly numerous. Yet in humans, mechanistic accounts linking species-relevant early life experiences to variability in cognitive developmental outcomes is scarce. Across species, environmental unpredictability shapes adaptive behavioral traits, from foraging and aggression (*3–6*) to altered stress physiology (*7*, *8*), to learning and executive control (*9*, *10*), suggesting that variability in early conditions can canalize different developmental solutions (*11*, *12*). Here we investigated the interactions between infant gut microbial composition and behavioral responses to maternal unpredictability to characterize mechanisms of phenotypic plasticity in early human development.

Humans are altricial, meaning that infants rely on caregivers for survival and learning (*13*). A substantial literature shows that caregivers differ in the predictability of behavioral sequences delivered to infants during daily interaction, and that this behavioral unpredictability is measurable from temporal transitions among caregiver sensory behaviors (*9*). High maternal unpredictability has been associated with poorer later learning, self-regulation, and inhibitory control in both animal models and human work (*9*, *14–18*). In previous work in the cohort used in this analysis (*19*), infants tested in two separate countries (Malawi, South Africa) dynamically adjusted their looking times based on moment-to-moment changes in their caregiver’s unpredictability (*20*), in a manner that is consistent with online optimization of attention in the service of learning (*21*). Indeed, in the same cohort, caregiver unpredictability in very early infancy was associated with reduced neural signatures of infants’ statistical learning months later (*10*). Thus, precisely how infants behaviorally respond to early life maternal unpredictability *in the moment,* and the mechanisms that enable this response, may provide critical insight into how maternal unpredictability might tune long-term developmental outcomes.

Thus, we focused this analysis on infants’ moment-to-moment behavior, operationalized as visual orienting behavior (VOB) shifts to and from the mother during naturalistic dyadic interactions. This choice is grounded in classic findings that infants rapidly extract regularities and adjust visual attention orienting behavior to statistical structure in their environment (*22–24*). Why might this impact later self-regulation and inhibitory control? Development proceeds along broad caudal to rostral cortical gradients: early-maturing sensory cortices such as vision can influence, and be integrated with, later-developing control systems, including prefrontal cortex (*25–27*). Thus, we expected that differences in early visual orienting behavior response to the mother, enabling or precluding learning in the moment, could plausibly be a mechanistic link between the observed association of early maternal unpredictability and later inhibitory control. Building on that rationale, we assessed whether infants exposed to more unpredictable maternal sequences adjust their visual orienting behavior patterns in ways that forecast later inhibitory control skills (*9*, *14*, *15*).

As noted, infant behavior responses in the moment, and later developmental trajectories, are rarely deterministic. As such, exposing biological drivers of variability is critical. Human infant studies that link longitudinal behavior with genetic, epigenetic, or microbiome composition are rare. The infant gut microbiome is dynamic in the first months of life and typically dominated by *Bifidobacterium* species whose relative balance varies across infants, with recurrent patterns across geographies (*28–30*). Microbial metabolites, including short-chain fatty acids (SCFAs), glutamate/GABA, and tryptophan-derived indoles/kynurenines, modulate microglia, excitation/inhibition (E/I) balance, and sensitive-period plasticity (*31–36*).

To probe mechanism rather than correlation alone, we paired naturalistic mother-infant interactions with shotgun metagenomics collected concurrently (±30 days) with behavior (Figure 1). We used metagenomics to resolve species-level contrasts central to early life (e.g., *Bifidobacterium breve* vs. *B. longum*) and to quantify gene families/pathways bearing on neuroactive chemistry (e.g., SCFA synthesis modules; tryptophan and glutamate/GABA pathways). Analytically, we (i) tested whether taxonomic/functional β-diversity explained variance in VOB (PERMANOVA on the Bray-Curtis dissimilarity landscape), (ii) identified species-VOB associations with compositional-aware transforms and FDR control, (iii) used Random Forests to assess multivariate predictive importance in a nonlinear setting, and (iv) performed feature-set enrichment to ask whether neuroactive gene sets are enriched or depleted as a function of maternal unpredictability, VOB, and their interaction. This strategy links a species-relevant environmental exposure (caregiving unpredictability) to an online behavioral visual orienting behavior policy and a biochemical substrate capable of supporting or constraining that 127 policy during early sensitive windows (*37–39*).

**Figure 1:**
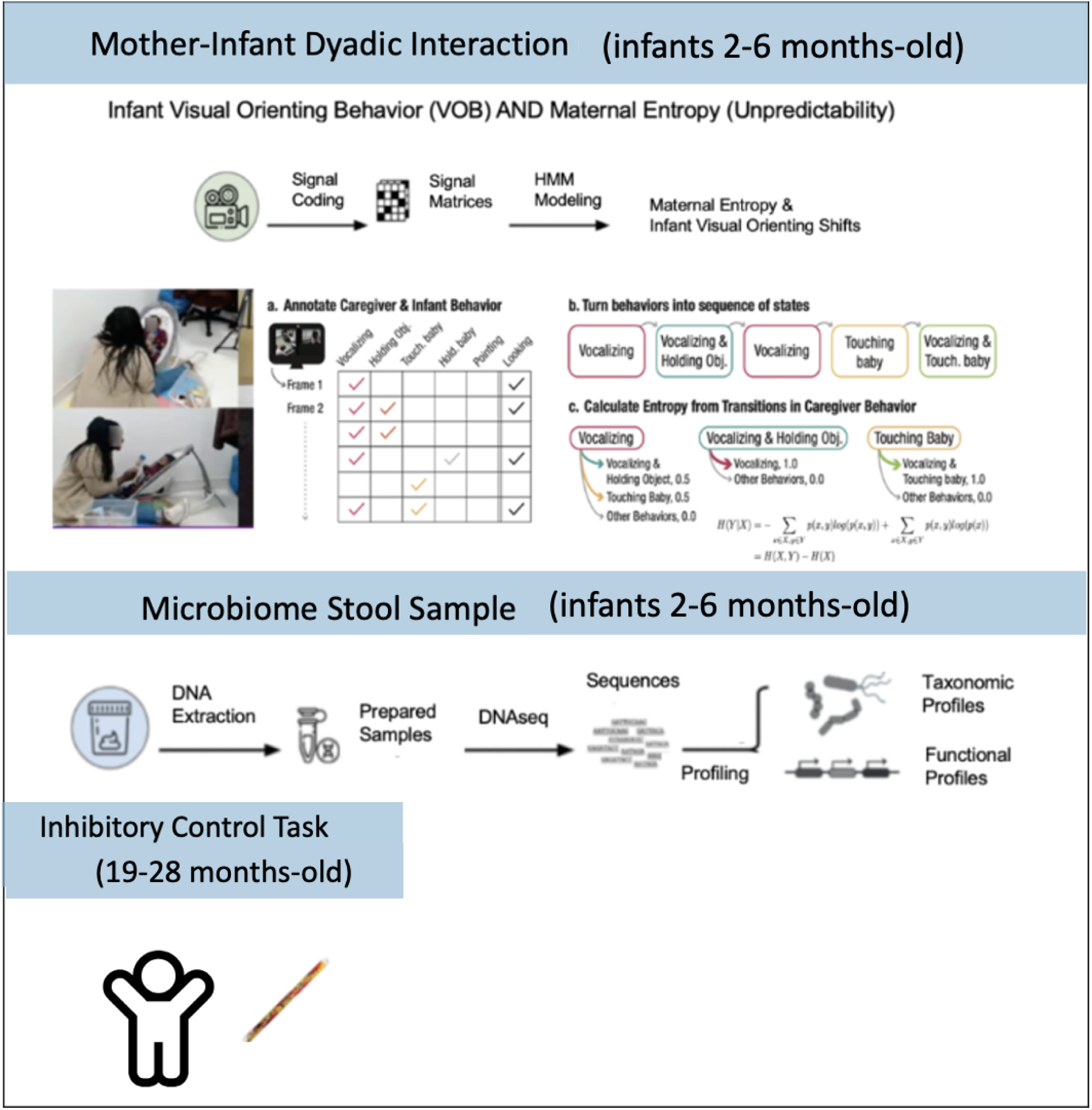
**Top Panel.** Participants engaged in a dyadic interaction when infants were between 2-6 months-old. Videos were hand-annotated for five maternal behaviors and simultaneous infant visual attention orienting behavior shifts (a). Caregiver unpredictability was calculated using behavioral transitions (b&c). **Middle Panel.** Microbiome stool samples were collected, prepared, and sequenced, resulting in analysis of functional and taxonomic profiles. **Left Panel.** Children completed an inhibitory control tasks when they were between 19-28 months.

## Results

### Unpredictable caregiving predicts a mature-for-age visual orienting behavior strategy

A total of 256 mother-infant dyadic interaction videos (118 female), each lasting approximately 5 minutes, were hand annotated for this analysis based on the coding scheme (*see Supplemental Materials*) used in (*9*, *40*). Three cameras captured the mother, the infant, and all together the dyad during a semi-naturalistic interaction (without toys or other objects) between a mother and her infant at a testing site in Cape Town, South Africa. These data are taken from a large cohort study with multiple assessments at longitudinal timepoints. In Table 1, we report the sample sizes for each assessment, as well as descriptive statistics for infant and mother demographic variables. In previous work using a subset of this sample’s data (*n* = 115 of the *N* = 256), paired with data from a sample of parent-infant dyads in Malawi, we showed that maternal unpredictability was associated with the total duration of infant looking behavior at the mother (*20*). The infant visual orienting behavior reported in this sample is correlated with, but is not the same as, the looking duration metric reported in the subset of South African infants tested in Forest and colleagues (*10*). Moreover, the inclusion of the inhibitory control and microbiome metrics is unique to this study.

**Table 1:**
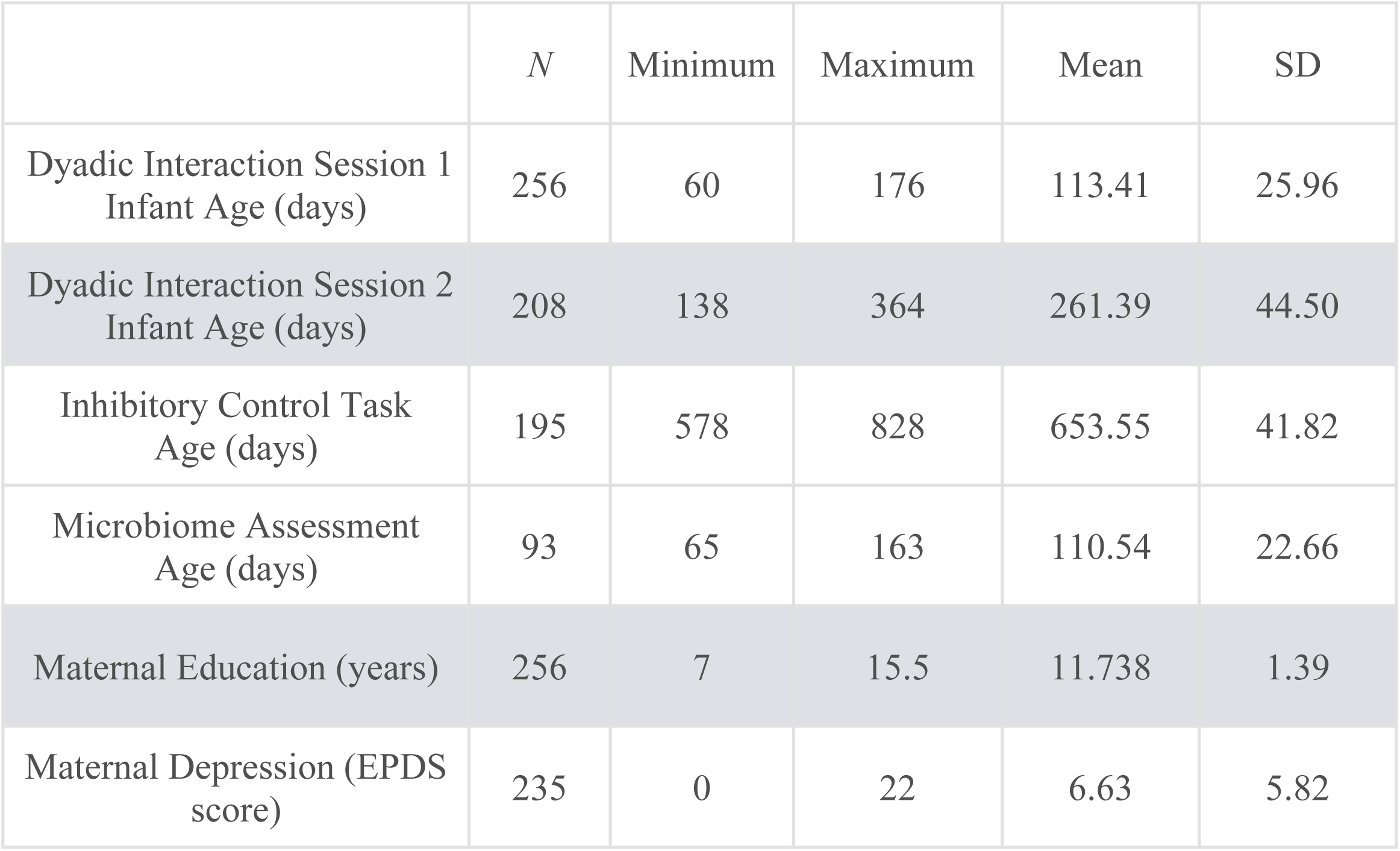
Sample Demographics.

|  | *N* | Minimum | Maximum | Mean | SD |
| --- | --- | --- | --- | --- | --- |
| Dyadic Interaction Session 1<br>Infant Age (days) | 256 | 60 | 176 | 113.41 | 25.96 |
| Dyadic Interaction Session 2<br>Infant Age (days) | 208 | 138 | 364 | 261.39 | 44.50 |
| Inhibitory Control Task<br>Age (days) | 195 | 578 | 828 | 653.55 | 41.82 |
| Microbiome Assessment<br>Age (days) | 93 | 65 | 163 | 110.54 | 22.66 |
| Maternal Education (years) | 256 | 7 | 15.5 | 11.738 | 1.39 |
| Maternal Depression (EPDS<br>score) | 235 | 0 | 22 | 6.63 | 5.82 |

We first examined the relationship between maternal unpredictability and shifts in infant visual orienting behavior. We defined VOB as the number of infant gaze shifts to or from the mother, the information source during the dyadic interaction. Based on prior work that examines infant looking behavior to unpredictable information (*21*), we reasoned that if maternal behavior is unpredictable, and therefore offers little learnable information, higher rates of shifting visual orienting to other possible sources of information may be a strategy for maximizing information uptake. However, visual orienting behavior shifts, to the extent that they reflect oculomotor control, can be slow to develop, with very early infancy (∼until 4-6 months) marked by rapid change and growth in oculomotor control (*26*). Therefore, showing higher rates of VOB shifts, in response to unpredictable maternal behavior, may reflect a mature-for-age pattern in infants in our 2-6 month-old infant sample.

Table 2 reports the correlations between maternal unpredictability, VOB, and maternal and family demographics. Only VOB shifts correlated with maternal unpredictability, such that higher maternal unpredictability was associated with more infant VOB shifts during dyadic interaction. To confirm that this correlation was not driven by infant age, we conducted an ANCOVA for the dependent variable VOB, with infant age (in days) as a covariate and maternal unpredictability as a continuous variable. VOB shifts increased with infant age, as would be expected based on the developmental visual orienting literature, *F*(1,253) = 6.510, *p* = 0.011, ηp² = 0.025. VOB shifts also independently increased with higher values of maternal unpredictability, while controlling for infant age in the model, *F*(1,253) = 29.189, *p* < 0.001, ηp² = 0.103.

**Table 2:** Maternal Unpredictability Correlations.

|  | Maternal Unpredictability | Infant VOB | Infant Age Session 1 | Maternal Education |
| --- | --- | --- | --- | --- |
| Infant VOB | .35** |  |  |  |
| Infant Age Session 1 | .22** | .22** |  |  |
| Maternal Education | .006 | -.06 | .05 |  |
| Maternal Depression (EPDS) | -.01 | .02 | -.05 | -.13* |

\* $p < .05$ , \*\* $p < .01$ , missing values replaced with series means

We next asked whether infants whose mothers were higher on the unpredictability continuum had a higher VOB *for their age* than those whose mothers were relatively more predictable. We extracted the residuals from a model that examined only the effects of infant age on VOB, *F*(1,254) = 13.066, *p* < 0.001, ηp² = 0.049, and correlated the resulting residual values with maternal unpredictability. As maternal unpredictability increased, so did the VOB *for age, r*(254) = 0.314, *p* < 0.001. Figure 2A shows higher age-adjusted VOB among infants whose mothers are more unpredictable.

**Figure 2:**
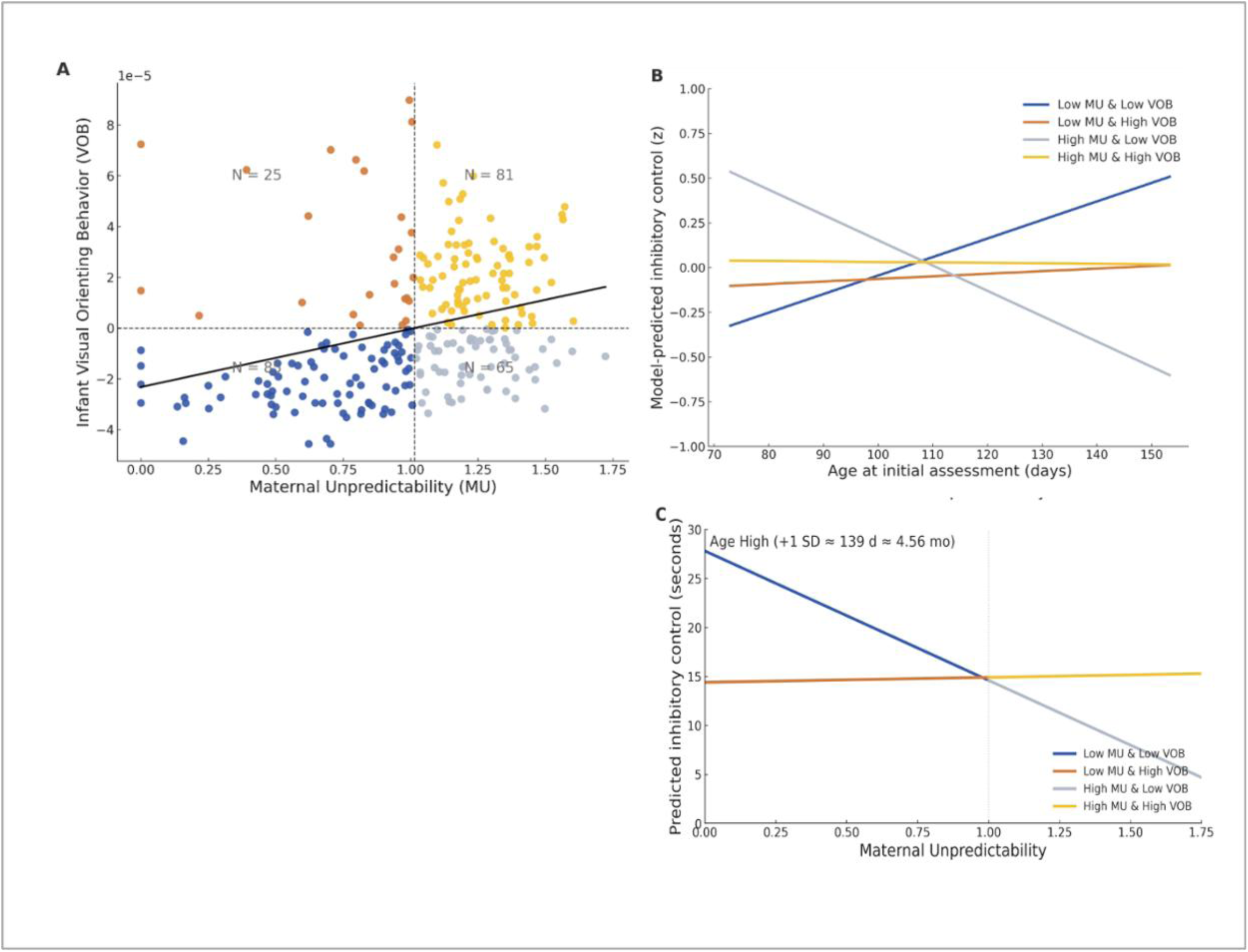
Maternal unpredictability, infant VOB, and later inhibitory control. **(A)** Illustrates the association between Maternal Unpredictability (MU) and infant visual orienting behavior (VOB) shifts (residual values for age) in 2-6 month-old infants. Higher VOB indicates higher values than expected for age. **(B)** Simple slopes illustrate the relationship between the four groups delineated in panel A and inhibitory control task performance when the same infants were 19-28 months-old. **(C)** illustrates the simple slopes for maternal unpredictability predicting inhibitory control separately when infants are tested in the upper end of the age range.

An examination of the data pattern in Figure 2A prompted further analyses: VOB shifts are not uniformly distributed across maternal unpredictability. Rather, low maternal unpredictability is associated with relatively slower-for-age VOB developmental patterns, or fewer VOB shifts, while increasingly higher unpredictability is associated with more variability in VOB responses. We color-coded each mother-infant dyad on the two variables of interest for illustration: infant VOB *for age* (mean split to create high/low VOB groupings) and unpredictability (mean split to create high/low maternal unpredictability groupings). Figure 2A shows that VOB shifts are not uniformly distributed across maternal unpredictability groupings, Pearson Chi-square (1, *N* = 256) = 27.74, *p* < 0.001 confirms group distributions differ statistically. The majority of infants whose mothers were low in unpredictability during the dyadic interaction had low values for VOB shifts for age (*n* = 85 vs. high VOB *n* = 25). In contrast, higher maternal unpredictability was associated with more variability in VOB shifts for age (*n* = 81 with higher VOB shifts and *n* = 65 in the lower quadrant), ostensibly reflecting variable behavioral strategies in response to a similar level of maternal unpredictability.

In order to verify persistence of effects, we tested the dyads using the same unstructured dyadic interaction protocol three months after their first session. See Table 1 for sample size retained and infant age details. First, there was a correlation between maternal unpredictability at the two timepoints, *r*(205)= 0.163, *p* = 0.019, indicating that maternal unpredictability is a general maternal characteristic rather than a one-time maternal state. Second, maternal unpredictability was again positively correlated with VOB, *r*(205)= 0.19, *p* = 0.006. These results suggest a persistence of effects into the second half of the first postnatal year. Finally, maternal unpredictability at the early testing time was not correlated with VOB at the later time, and later maternal unpredictability was not correlated with VOB at the earlier testing time (all *p* > 0.258). Taken together, these results demonstrate that variability in VOB in response to maternal unpredictability is best explained as a visual orienting strategy used during moment-to-moment dyadic interactions to manage learnability of input, rather than a driver of general disorganized visual attention orienting development in infancy.

### High infant VOB in the context of high maternal unpredictability is associated with resilience in later inhibitory control scores

Davis and colleagues (*41*) showed that high maternal unpredictability in infancy was associated with poor effortful control in childhood. First, we asked whether we would replicate, in this South African cohort, the negative relationship obtained with control and maternal unpredictability in Western cohorts. A subset of infants performed an inhibitory control task when they were between 19-28 months of age (*n* = 195, 90 female). Participants were presented with a shiny, glittery toy and given the rule that they must wait (up to 30 seconds) before approaching the toy for play. The inhibitory control score is the z-scored wait time in seconds. This score was not correlated with inhibitory control task test age, *r*(193) = 0.059, *p* = 0.409).

We fit a general linear model predicting inhibitory control from maternal unpredictability, infant VOB, and infant age at initial assessment (and their interactions), with infant sex and maternal depression (EPDS) as covariates. There was no significant main effect of sex, *F*(1, 185) = 0.677, *p* = 0.412, ηp² = 0.004. We observed trend-level main effects of infant age, *F*(1, 185) = 3.524, *p* = 0.062, ηp² = 0.019, and VOB, *F*(1, 185) = 3.338, *p* = 0.069, ηp² = 0.018, and a significant main effect of maternal unpredictability, *F*(1,185) = 4.006, *p* < 0.05, ηp² = 0.021. Interactions were as follows: age by maternal unpredictability, *F*(1, 185) = 4.381, *p* < 0.05, ηp² = 0.023; age by VOB, *F*(1, 185) = 3.163, *p* = 0.077, ηp² = 0.017; maternal unpredictability by VOB, *F*(1,185) = 4.112, *p* < 0.05, ηp² = 0.022; and an age by maternal unpredictability by VOB three-way interaction, *F*(1, 185) = 4.092, *p* < 0.05, ηp² = 0.022. Higher maternal unpredictability was associated with lower inhibitory control scores. The interaction with age indicates that assessments conducted in infants that were older in the 2–6 month age range were more likely to reveal this effect.

To interpret the patterns revealed by the maternal unpredictability by age by VOB three-way interaction, we examined the simple slope of maternal unpredictability at ±1 SD of infant age and VOB (covariates fixed). At the higher end of the 2–6 month age range (+1 SD ≈ 138.9 days ≈ 4.56 months), higher maternal unpredictability in the context of low infant VOB (grey markers in Figure 2A) predicted shorter wait times (poorer inhibitory control), *b* = −1.431, *SE* = 0.437, *p* = 0.001 (HC3 *p* = 0.006). In contrast, when VOB was high (+1 SD; yellow markers in Figure 2A), the maternal unpredictability slope was indistinguishable from zero: *b* = +0.589, *SE* = 0.547, *p* = 0.287 (HC3 *p* = 0.304). The difference between the two slopes (Low − High VOB) was sizable (*Δb* = −1.064, *SE* = 0.713). At the younger end (−1 SD ≈ 88.0 days ≈ 2.89 months), maternal unpredictability slopes were not significant whether VOB was low (*b* = −0.233, *SE* = 0.383, *p* = 0.545) or high (*b* = +0.269, *SE* = 0.269, *p* = 0.318). This pattern suggests that higher infant VOB attenuates the negative association between maternal unpredictability and later inhibitory control, particularly at older infant ages. We next explored potential mechanisms underlying variability in VOB responses in early infancy.

### Microbiome composition is associated with the VOB strategy

We examined microbiome data for evidence of association between microbial composition and VOB. Data on both the gut microbiome and maternal-infant interactions were available for *n* = 93 (38 Female) of the same infants whose data are discussed above. Stool samples (*n* = 93, one per infant) were collected for metagenomic analysis concurrently (± 30 days) to PCI measurements. We used these data to analyze the effect of various factors on gut microbial community diversity at the taxonomic and functional gene levels. Specifically, we performed both taxonomic and functional profiling of the metagenomes using a uniform computational pipeline (see Materials and Methods). Taxonomic profiles consisted of species-level relative abundances (with an average 34 [*SD* = 14] species per sample). Functional profiles consisted of relative abundances of gene families and pathways (with an average 274,300 [*SD* = 110,700] gene functions per sample).

We began with a permutational analysis of variance (PERMANOVA), intended to preliminarily assess whether specific metadata could explain variance in the inter-sample diversity (β-diversity) of gut metagenomes. β-diversity was computed as pairwise Bray-Curtis dissimilarities between metagenomes. While factors such as age at collection, infant sex, maternal education, and maternal unpredictability did not significantly explain variance in community diversity (*p* > 0.05), VOB was significant for both taxa and genes (*R*² = 0.038, *p* = 0.003 for taxa; *R*² = 0.021, *p* = 0.012 for genes; Figure 3A). These data provided an initial indication that infants with high VOB may have a distinct gut-microbial signature.

**Figure 3.**
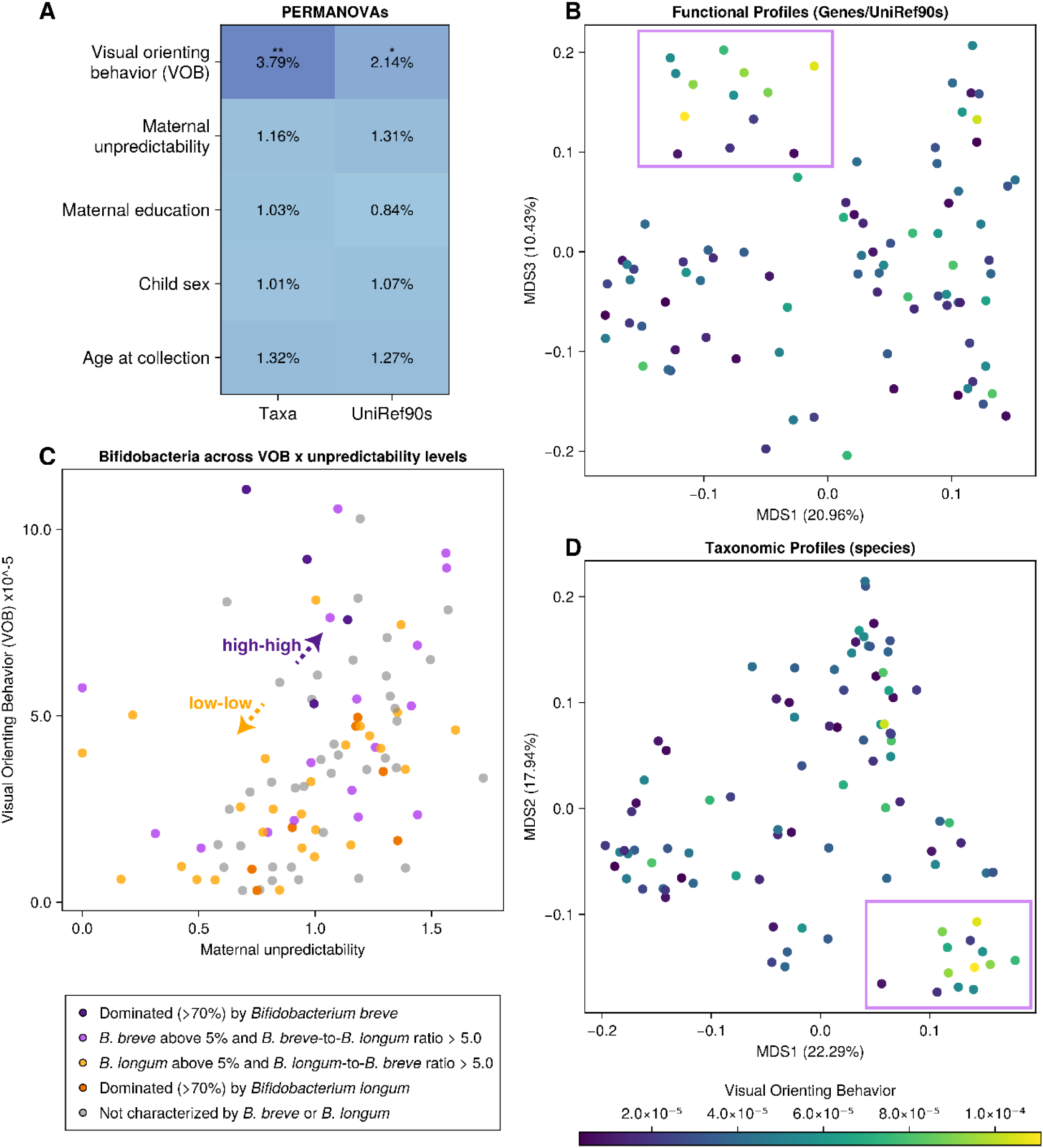
Infant Visual Orienting Behaviour (VOB) and microbial community profiles share variance explained at both taxonomic and functional levels. **(A)** Permutational analysis of variance (PERMANOVA) of taxonomic (species-level relative abundances) and functional profiles (UniRef90 gene function relative abundances) against the metadata of interest. Variations in these profiles significantly explain the variation in infant visual orientation behaviour. Markers above the number indicate significance level (* - p < .05; ** - p < .01; *** - p < .001). **(B)** Relationship between maternal unpredictability and infant visual orienting behaviour, with participants colored by taxonomic alpha diversity (a measure of within-sample diversity) obtained by calculating the Shannon diversity index. **(C)** Principal coordinate analysis (PCoA) by multidimensional scaling (MDS) of Bray-Curtis dissimilarities of functional profiles (Uniref90 gene functions). The percent variance explained by each Principal Coordinate is indicated on the X and Y axes. Samples (dots) are colored *a posteriori* by infant visual attention shifts. The high-attention cluster mentioned in the results is highlighted. **(D)** Principal coordinate analysis (PCoA) by multidimensional scaling (MDS) of Bray-Curtis dissimilarities of taxonomic profiles (species). The high-attention region mentioned in the results is highlighted.

To test this hypothesis directly, we conducted an ordination analysis on the β-diversity matrix of functional gene profiles using multidimensional scaling (MDS). This approach allows us to reduce complex, high-dimensional diversity patterns into a few interpretable axes that capture the major sources of variation across samples. The first three principal coordinates explained 45.65% of the variance. While the primary projection of PC1 vs. PC2 showed no appreciable patterns (Supplementary Figure S1), the projection of PC1 vs. PC3 (Figure 3B) revealed that children with higher VOB shifts form a visually distinct cluster.

To formally assess these clustering patterns, we applied k-means clustering. We selected the optimal number of clusters (*k* = 4) based on a combination of Within-Cluster Sum of Squares (WCSS) and Silhouette Score (Supplementary Figure S2A). Consistent with the visual separation in Figure 2B, one cluster stood out with a mean infant VOB nearly twice as high as the others. While most clusters had mean VOB values ranging from 3.15 × 10⁻⁵ (*SD* = 2.09 × 10⁻⁵) to 3.87 × 10⁻⁵ (*SD* = 2.54 × 10⁻⁵), this group exhibited a significantly higher mean of 6.13 × 10⁻⁵ (*SD* = 3.58 × 10⁻⁵, *p* < 0.001; Supplementary Figure S2 C-D). The combination of a clear microbiome-derived cluster of infants with high VOB shifts and our earlier observation that infants with high VOB are almost twice as likely to have high unpredictability mothers (*n* = 81) than low unpredictability mothers (*n* = 25) (Figure 2A), suggests that infants with greater visual orienting behavioral shifts exhibit unique microbial diversity profiles at the functional gene level.

To determine whether taxonomic β-diversity patterns would mimic the VOB distribution patterns found in the functional gene data, we replicated the MDS ordination analysis using species-level data. The first three principal components explained 49.53% of the variance in taxonomic-level βdiversity (Figure 3D). While the projected sample scores did not exhibit the same distinct structure, the principal coordinates revealed that a small subset of microorganisms drove variation in community diversity. PC1 (22.28% of total variance) was primarily associated with typically exclusive infant enterotypes dominated by either *Bifidobacterium longum* (*R*² = 0.825) or *Bifidobacterium breve* (R² = 0.347). Notably, PC1 was significantly correlated with VOB (*R*² = 0.116), despite VOB not being included as a factor in the ordination analysis. PC2 (17.93% of total variance) was associated with a balance between prominent *Bifidobacterium* spp. (*B. longum*, R² = 0.090; *B. breve*, R² = 0.347; *Bifidobacterium bifidum*, *R*² = 0.136) and other genera commonly found in the developing guts of infants (*Ruminococcus gnavus*, *R*² = 0.315; *Erysipelatoclostridium ramosum*, *R*² = 0.153; *Escherichia coli*, *R*² = 0.175; Supplementary Figure S3). We followed this result with exploratory data analysis between the exclusive species of *Bifidobacterium* (*B. breve* and *B. longum*) and their relationship to the non-uniform distribution of VOB with the across different maternal unpredictability levels. Although initial visualizations suggested that the balance between *B. breve* and *B. longum* was associated with the topology of the VOB-Entropy space, with the high-high quadrant characterized by a predominance of *B. breve*, and the low-low quadrant by a predominance of *B. longum* (Figure 3C), no significant results were found when MANOVA was conducted between the VOB-Unpredictability space and the data quadrant assignments of each sample.

Building on the ordination patterns and exploratory data analysis, we tested individual species for associations with maternal unpredictability and VOB. We modeled z-normalized VOB as an outcome for each microbial feature (*m* = 45) that met a minimum prevalence of 5% and a mean relative abundance ≥ 1%, while controlling for age at stool collection. Microbial relative abundances were arcsine-transformed. No microbial species showed a significant association with maternal unpredictability after false-discovery rate (FDR) correction using the BenjaminiHochberg method. However, for VOB, *B. longum* (β = -0.117, *p* = 0.003, *q* < 0.20) and *B. breve* (β = 0.109, *p* = 0.007, *q* < 0.2) remained significant after FDR correction (Figure 4A-B). This is consistent with the taxonomic profile ordination findings, where those species explain variance on PC1, which, in turn, is independently correlated with VOB.

**Figure 4.**
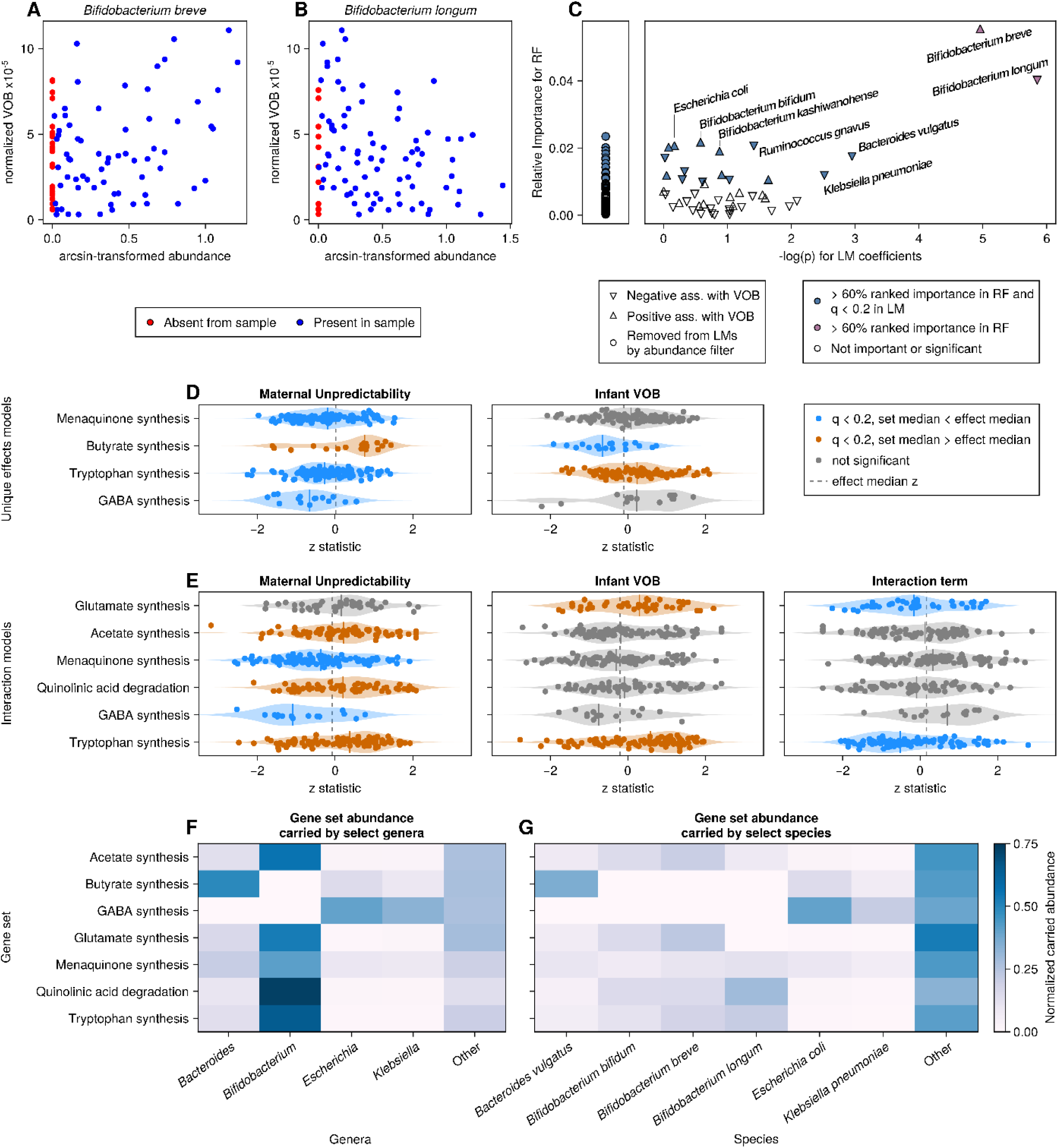
Microbial gene functions and their carrier species are associated with infant VOB. (A/B) Znormalized VOB as a function of arcsin-normalized species abundance for both FDR-passing microbes, *B. breve* (A) and *B. longum* (B). Colors indicate whether the microbe was detected in the profiled metagenome. (C) Comparison between the Random Forest feature importance (measured as Mean Decrease in Impurity - MDI) and unique effect significance (p-value) for each input species towards the prediction of infant VOB. Main panel marker shapes indicate whether species were positively or negatively associated with the outcome on linear models. The side panel shows species that were removed from linear models by the abundance filter, which only have an RF importance associated (D-E) Feature set enrichment analysis (FSEA) of microbial neuroactive genes and maternal unpredictability, infant visual attention and their interaction. (D) shows results for Model 1 (see Methods) accounting for unique effects only, while (E) shows results for Model 2, accounting for interaction between the factors. FSEA results are shown for neuroactive gene sets where at least one tested component, maternal unpredictability, infant visual attention, or the interaction term between the maternal unpredictability and infant visual attention, had a significant hit (q < 0.2). Dots indicate the Z-statistic from logistic regression for each gene in a gene set; vertical bars indicate the median Z-statistics for the whole gene set. (F-G) Proportion map of carrier for each characterized gene function on each neuroactive gene set shown on Panels D-E. (F) Gene set abundance carried by select genera; (G) Gene set abundance carried by select species. Shade strength indicates the proportional count of detected genes, in each gene set, carried by taxa in columns.

While individual associations provide insights into effect sizes and significance, gut microbial community effects should be considered within their broader compositional context. To address this, we used a nonparametric multivariable model, Random Forest (RF), to predict VOB from all microbial features that met the 5% prevalence filter, regardless of mean abundance (*m* = 149), again controlling for age. In a three-fold-cross-validation setting, the model achieved a Root Mean Square Error (RMSE) of 1.10 and a Pearson correlation of 0.231 between the predicted and ground-truth z-normalized VOB in validation sets. We then extracted the variable importance ranking from the RF model and compared it to the significance of effects from linear models. As expected, both *B. breve* and *B. longum* ranked highest in both scales, even surpassing the infant age at sample collection feature (Figure 4C). Additionally, *Klebsiella pneumoniae* achieved high ranks for both importance and effect size significance. A third group, including *Bacteroides vulgatus*, *Enterococcus faecalis*, and *E. coli*, showed high importance in RF models despite showing modest effects in linear models. Additionally, we probed whether microbial features would improve the prediction of VOB alongside maternal unpredictability. To do so, we built a second RF model predicting VOB from a combination of Maternal Unpredictability and 30 top-ranking microbial features. Predictions achieved a Pearson R of 0.407 with the ground truth, which, despite being a numerical improvement over the prior correlation between VOB and Maternal unpredictability (*R = 0*.367), was not statistically significant with the sample size (*p* > 0.05, *n* = 93).

### Neuroactive metabolic potential is context-dependent

Building on the taxonomic-level analysis that revealed significant effects of gut microbes in infant VOB, we sought to determine whether the variations in species composition also reflected functional shifts in microbial communities, particularly in relation to neuroactive gene pathways. We hypothesized that microbial gene functions associated with neuroactive metabolites could exhibit enrichment or depletion patterns based on developing levels of VOB and maternal unpredictability due to their correlated topology. To test this hypothesis, we conducted feature set enrichment analysis (FSEA) to assess the relationship between maternal unpredictability, infant VOB, and the presence of specific microbial genes. The objective of this analysis was to determine whether different sets of genes are enriched or depleted at different levels of phenotypes of interest.

We employed logistic regression models to examine the association between gene presence/absence and these behavioral outcomes, while adjusting for control variables (age and sex). Several gene sets involved in the metabolism of neuroactive molecules (*36*) were significantly enriched or depleted in relation to maternal unpredictability and VOB. Specifically, when considering only the unique effects of these two measures (Methods, Equation 1), gene sets involved in tryptophan synthesis were significantly enriched (ES = +0.133, *q* = 0.144) in children with higher VOB, while genes involved in butyrate synthesis were significantly depleted (ES = -0.308, *q* = 0.144). For maternal unpredictability, the gene sets for GABA synthesis (ES =-0.392, *q* = 0.047), tryptophan synthesis (ES = -0.185, *q* = 0.148), butyrate synthesis (ES = +0.388, *q* = 0.165), and menaquinone synthesis (ES = -0.140, *q* = 0.165) also showed significant enrichments or depletions (Figure 4D). Notably, for the gene sets that showed significance for both terms, the sign of enrichment scores was always opposite, which indicates that, independent of each other, gene sets characteristic of high values of VOB are different from those characteristic of high values of maternal unpredictability.

When the model was expanded to include an interaction term between maternal unpredictability and VOB (Figure 4E, Equation 2), the enrichment landscape changed to include glutamate synthesis as a significant enrichment for VOB (ES = +0.211, *q* = 0.138) and a depletion for the interaction term (ES = -0.266, *q* = 0.165). Tryptophan synthesis remained strongly significant across the board, on both effects and interaction term (ES = +0.171, *q* = .058 for maternal unpredictability; ES = +0.291, *q* = 7.1 × 10^-6^ for VOB; ES = -0.238, *q* = 5.7 × 10^-5^ for the interaction). For maternal unpredictability, GABA synthesis and menaquinone synthesis were significantly depleted (respectively, ES = -0.342, *q* = 0.07; ES = -0.138, *q* = 0.150) while acetate synthesis and quinolinic acid degradation were significantly enriched (respectively, ES = +0.138, *q* = 0.183; ES = +0.141, *q* = 0.48) in children with more unpredictable mothers.

The interaction model revealed that the combined effect of maternal unpredictability and infant VOB followed a distinct pattern, diverging from their individual contributions. Specifically, the interaction term consistently exhibited the opposite sign compared to the unique effects of maternal unpredictability and VOB, suggesting that microbial gene enrichment is not merely a linear function of either factor. Instead, this pattern indicates that the microbiome’s functional potential, particularly its role in neuroactive metabolism, modulates behavioral plasticity in a nonlinear way, shaped by both VOB and environmental unpredictability. Rather than maternal unpredictability directly influencing microbial function, these results suggest that microbial metabolic activity plays an active role in shaping behavioral variability, with distinct gene-functional signatures emerging in response to different combinations of VOB and unpredictability. This supports the broader idea that gut microbial composition and function influence neurodevelopmental processes by modulating the biochemical landscape underlying behavioral flexibility.

To further connect these functional shifts to specific microbial taxa, we examined the distribution of neuroactive gene sets across the key genera identified in our taxonomic analyses. Unsurprisingly, neuroactive gene sets were predominantly carried by commonplace commensals of the infant gut that were highlighted in the previous analyses: *Bifidobacterium*, *Bacteroides*, *Klebsiella*, *Veillonella*, and *Escherichia* (Figure 4E). Among these genera, *Bifidobacterium* is an especially predominant carrier of neuroactive gene sets, especially those that were more significantly associated with VOB, such as tryptophan and glutamate synthesis (Figure 4F). Interestingly, GABA and butyrate synthesis are not carried at all by bifidobacteria, being instead split between other genera that were previously implicated in the nonlinear prediction of VOB (Figure 4C). We followed that analysis by a breakdown of species among those main carrier genera to look for differences in the carried repertoire between the different species of *Bifidobacterium*. We discovered that, although *B. longum* and *B. breve* are the topmost abundant *Bifidobacterium*, their functional repertoire differs in terms of neuroactive gene sets carried. Notably, *B. longum*, who carries the most abundance of tryptophan synthesis and quinolinic acid degradation, did not contribute at all to glutamate synthesis, which was in turn attributed to *B. breve* and other less prevalent species such as *B. bifidum* (Figure 4G). This finding provided a mechanistic link between microbial composition and neuroactive metabolic plasticity, reinforcing the idea that species-level differences in the microbiome do not merely correlate with behavior but actively contribute to its modulation in the context of environmental unpredictability, via the metabolic framework enabled by genes carried by different microbial species.

## Discussion

Our findings support a model in which environmental unpredictability, here measured with maternal behavior, calibrates infant behavioral plasticity, measured here in visual orienting shifts response, with microbial community composition shaping the biochemical capacity for infant behavior via neuroactive metabolic pathways. In naturalistic parent-infant interactions, higher maternal unpredictability was associated with more frequent infant visual orienting behavior shifts, a mature-for-age strategy (Figure 2A). Longitudinally, greater maternal unpredictability in early infancy predicted poorer inhibitory control at follow-up. While VOB did not fully explain the relationship between early maternal unpredictability and later inhibitory control scores, higher VOB attenuated the relationship between high maternal unpredictability and later inhibitory control. As shown in Figure 2B, infants exposed to high maternal unpredictability but who exhibited elevated VOB displayed inhibitory control outcomes more comparable to those from low-unpredictability environments with similar VOB, and better than infants in high unpredictability environments but with low VOB.

To summarize, in predictable contexts, a stability-oriented behavioral response (fewer visual orienting shifts) appears as an adaptive, and indeed highly prevalent strategy (∼77% of infants showed this pattern), with positive later impact on inhibitory control scores (Figure 2B, blue line). In contrast, in the context of high maternal unpredictability, where the learning signal is ostensibly diminished, infants who shifted gaze frequently, a mature-for-age strategy (Figure 2B, yellow markers), may optimize information uptake. This high VOB pattern was observed in a small subset of the sample (41% overall, and ∼55% in the context of high maternal unpredictability). In the context of unpredictable caregivers only, this higher VOB strategy was associated with a more resilient later inhibitory control profile (Figure 2B, yellow versus grey lines). We interpret this as evidence of phenotypic plasticity. What enables some infants to engage in high VOB for age in the context of maternal unpredictability (*n* = 81, Figure 2A), while a substantial subset (*n* = 65) of infants who experience high maternal unpredictability engaged in a less adaptive but more overall common strategy?

The microbiome converged on a *Bifidobacterium* sp. gradient that tracked VOB in early infancy: higher VOB, which was more than 3x as likely in high maternal unpredictability contexts (Figure 2A, yellow vs orange markers), aligned with a shift toward *B. breve* and away from *B. longum* (Figure 3). Random-forest rankings reinforced the centrality of *Bifidobacterium* species, with *Klebsiella pneumoniae* and *Bacteroides vulgatus* emerging as secondary signals. Both *B. breve* and *B. longum* are dominant early colonizers that produce neuroactive metabolites including short chain fatty acids (SCFAs), glutamate, and tryptophan-derived products that are implicated in neurotransmission and neurodevelopment (*30*, *31*, *35*, *37–39*, *42*). SCFAs, especially acetate from *Bifidobacterium* sp. (here *B. breve*, Figure 4G) regulate microglia and synaptic pruning essential for critical-period plasticity (*34*, *43*). Microbial glutamate production and conversion to GABA modulate E/I balance in developing visual cortex (*37*, *38*), and microbial tryptophan metabolism yields serotonin and kynurenine-pathway metabolites that stabilize circuits and regulate plasticity (*44*). We interpret our findings in the context of this literature.

Bacterial community composition was not sufficient to explain *adaptive* infant responses (low VOB in low maternal unpredictability contexts and high VOB in high maternal unpredictability contexts). The critical question is what enabled high VOB for age in high maternal unpredictability contexts, especially given the overall prevalence of low VOB during dyadic interaction in our age range? Pathway enrichment analyses showed that higher VOB aligned positively with tryptophan-synthesis enrichment and butyrate synthesis depletion (an SCFA) and, in the interaction models, glutamate synthesis enrichment. Critically, this pattern was not consistent across maternal unpredictability. The maternal unpredictability by VOB interaction term was *negative* for tryptophan and glutamate synthesis (Figure 4E) indicating that the positive coupling between VOB and glutamate and tryptophan *attenuates* as unpredictability increases. Instead, the results point to alternative pathway profiles in conditions of high maternal unpredictability.

Comparing the unique-effects model to the interaction-inclusive model clarifies these patterns. In the unique-effects model (Figure 4D), maternal unpredictability showed depletions for GABA and tryptophan synthesis and no acetate/quinolinic signal, whereas adding VOB and the maternal unpredictability by VOB term (Figure 4E) left the GABA and menaquinone results for maternal unpredictability unchanged, reversed maternal unpredictability’s association with tryptophan to enrichment, and newly revealed maternal unpredictability-linked enrichment of acetate synthesis and quinolinic-acid degradation. At the same time, VOB remained positively aligned with tryptophan (and, in the interaction model, glutamate), but the maternal unpredictability by VOB interactions were negative for tryptophan and glutamate, again indicating that the VOB to Trp/Glu coupling attenuates as maternal unpredictability increases (Figure 4E). Taken together, under high maternal unpredictability, high VOB coincides with an alternative pathway portfolio that includes signals for SCFAs (acetate) and quinolinic handling, with contributions consistent with species non-redundancy (*B. breve vs B. longum*) and additional taxa (e.g., *K. pneumoniae*, *B. vulgatus*; Figure 4C, 4F–G).

Infant-type *Bifidobacterium* sp. cannot fully synthesize tryptophan *de novo* but convert it to indole metabolites (ILA/IAA/I3CA) that activate AhR/Nrf2 and temper inflammation, indirectly affecting tryptophan availability and signaling (*45*, *46*). In more inflamed contexts, host IDO/TDO shunt tryptophan toward kynurenine to quinolinic acid, reducing plasticity-supporting chemistry and increasing stress-linked metabolites (*47*), patterns reported under high unpredictability (*16*, *48*, *49*). In our data, maternal unpredictability was associated with enrichment of acetate synthesis and enrichment of quinolinic-acid degradation when VOB and the maternal unpredictability by VOB term were included (Figure 4E), suggesting a shift under high maternal unpredictability toward an SCFA-supported profile. Acetate, predominant in early infancy and a major *Bifidobacterium* product, can fuel colonocytes, reinforce the barrier, and modulate immune/microglial tone, offering a plausible biochemical scaffold that could help sustain high VOB despite environmental volatility. By contrast, quinolinic acid is a pro-inflammatory, NMDA-agonist kynurenine metabolite; enrichment of its degradation genes is consistent with increased capacity to catabolize/neutralize quinolinic acid under high maternal unpredictability. Together, these findings point to re-routing of tryptophan fate and support in high maternal unpredictability, away from further up-coupling of tryptophan/glutamate and toward acetate-centered support plus quinolinic handling, with species specialization (*B. breve* vs. *B. longum*; Figure 4F–G) and additional contributors (e.g., *K. pneumoniae*, *B. vulgatus*; Figure 4C) shaping the available pathway mix. As these results reflect gene carriage, confirming pathway activity will require metabolomic/transcriptomic and inflammatory readouts.

Conceptually, the pattern aligns with developmental-plasticity models wherein organisms balance flexibility and stability in response to environmental cues (*12*), is consonant with rodent data linking unpredictable maternal input to accelerated inhibitory maturation (*50*), and extends that framework by implicating microbial tryptophan/indole metabolism as a plausible route from environmental signals to phenotypic plasticity in human infants. In this view, maternal unpredictability sets the environmental challenge, VOB implements the adaptive strategy, and the microbiome supports or constrains that strategy via neuroactive metabolites. Rather than a oneway path from maternal behavior to microbiome, the data suggest a mediating role for infant behavioral plasticity, with microbial composition reflecting the strategy infants adopt under their environmental conditions. By shaping the biochemical space of neural plasticity, the microbiome may enable resilience that helps infants sustain adaptive function under variable conditions.

## Methods

### Cohort Study Participants Data Collection

The reported data are collected as part of a multi-site larger longitudinal cohort study designed to characterize the developmental trajectory of executive functions and its precursors from 0-1000 days (*19*). The relevant study population was recruited through clinics in Gugulethu, South Africa (*N* = 393). All protocols were approved by the University of Cape Town Health Sciences Faculty Research Ethics Committee (666/2021). Mothers signed informed consents on behalf of themselves and their infants. All consents and procedures were translated/communicated in the preferred family language, i.e., Xhosa in South Africa. This is an ongoing study. Infants are tested in dedicated laboratory spaces over 5 time-points, beginning when infants were between 2-6 months, and roughly every 6 months after. The first testing session battery included parent-infant interactions, baseline EEG, a VEP assessment, collection of stool and blood samples, and structural

MRI. The data reported here are shared and may be used in other publications that analyze the relationship of these variables with other collected constructs and measurements in the cohort to address novel questions. The data will not be reused to ask the same empirical questions addressed here.

### Subsample Participants and Power

Of the total infants tested, 393 infants ages 2 to 6 months provided full parent-infant interaction data. Of these, videos from 263 participants were randomly-selected for hand-annotation. Four were then excluded after determining that the infants were wearing MRI ear protection during the session and were unlikely to hear the mother clearly. Three additional sessions were excluded because only one condition (see below) was recorded. In all cases, the parent in the session was the mother and the infant’s primary caregiver. The final sample size of *N* = 256 (118 female) is larger than would be needed to detect a medium sized effect (0.25) at 95% power, as determined by a G*Power analysis (*N* = 164).

### Recording Apparatus and Parent-/Infant Interaction Protocol

Three tripods held 3 Logitech Webcams that connected to a computer loaded with ManyCam software. ManyCam is a software that temporarily synchronizes audio and video from multiple cameras. Mothers were seated on the floor facing infants, who were seated in car seats. One camera directly faced the infant, one faced the mother, and the final camera captured a side view of the dyad.

Mothers were instructed to play with the infant for 10 minutes in the same way they would at home. These 10 minutes were divided into two segments, one with culturally appropriate toys available (Toy condition) and another without them (No Toy condition). The order of these conditions was randomized so as to counterbalance which condition came first. Only the No Toy condition was hand annotated in the majority of participants, and those data are presented here. In previous work, we found a high correlation between maternal predictability in the Toy and No Toy conditions (*20*).

### Coding of Maternal Sensory Signals and Infant Visual Orienting Behavior

Maternal behaviors that are tactile, auditory, and visual sensory signals were coded following a coding manual developed by Davis and colleagues (*9*). This coding manual captures maternal sensory signals to her infant and accounts for infant eye-gaze in the context of a dyadic interaction continuously in real time. Behaviors were coded using Datavyu, a video coding and data visualization software that enables the microanalysis of behavior in dyadic interactions. Due to variability in duration of study sessions, total time of video-recordings ranged from x-y minutes. Prior to coding, video quality was assessed by evaluating camera angles, image resolution and audio clarity. In four separate passes, trained coders noted temporal onset and offset of behaviors listed in Table X under the following sensory categories: (1) Maternal Auditory, (2) Maternal Tactile, (3) Maternal Visual, and (4) Infant Visual. Maternal auditory signals were coded as mother vocalizing. Maternal tactile signals were coded as mother holding baby and mother touching baby. Maternal visual signals were coded as mother holds object and mother points. Each pass involved coding only a specified category of behaviors (ex. maternal verbal behaviors were coded during one pass, and in a completely separate pass, infant eye-gaze was coded) meaning all codes are entirely mutually exclusive. The onset of a behavior is defined as the first frame where the behavior occurs and offset as the last frame where the behavior occurs. Therefore, a behavior is coded for both its occurrence and duration. Identically categorized behaviors separated by less than 500 milliseconds were identified as a single event and behaviors separated by 500 milliseconds or more or categorized differently were annotated as separate events. Instances where mother and/or infant behaviors were not codable because they were not clear in the video or audio (i.e instances where mother went out of view to grab a new toy; instances where experimenter vocal input hinders clarity on what mother is saying) were also annotated and classified as unknown. This coding approach allows for the dyadic interactions and relationship of behaviors to be explored solely on the basis of co-occurrence in real time and not coder interpretation of contingency.

Each 10-minute coded interval underwent the following process: (1) primary coding (2) reliability coding, where a different, secondary coder codes the same ∼10-minute interval in a subset of videos and (3) reliability check, where a script computes the percent reliability of each column. If interobserver reliability was below 80%, the indicated coders revised their coding separately for that participant and the reliability check process was repeated until interrater reliability criteria of at least 0.8 were met for every maternal behavior.

### Measuring Maternal Unpredictability/Entropy & Infant Visual Attention Orienting Behavior

The coding process resulted in data points displayed as cells, each representing an interval in time. If a behavior was coded as “active” during the given interval, the corresponding cell was assigned a value of 1; otherwise, cells were filled in with 0s. For each participant, data include temporally aligned columns representing the onset and offset of each occurrence of any of the 5 maternal behaviors (vocalizing, holding baby, touching baby, holding object, pointing), as well as for infant gaze towards the mother or the object mother is in contact with. Because multiple maternal behaviors can happen simultaneously, each time a new instance of any of the five targeted behaviors is coded as active, a new behavioral state is generated, with the new state’s onset being the previous state’s offset. We operationalize discrete independent behavioral states as any combination of the 5 maternal behaviors. For example, a mother might vocalize and hold an object at the same time.

Broadly, Shannon Entropy calculations involve the Markov property, where the predictability of any one maternal behavioral state (n+1) is conditioned only on the immediately previous state (n). If we define X, Y as two discrete random variables, X being the initial state and Y being the final state of any two-state transitions in a stochastic process, we have the conditional entropy formula of Y given X as:

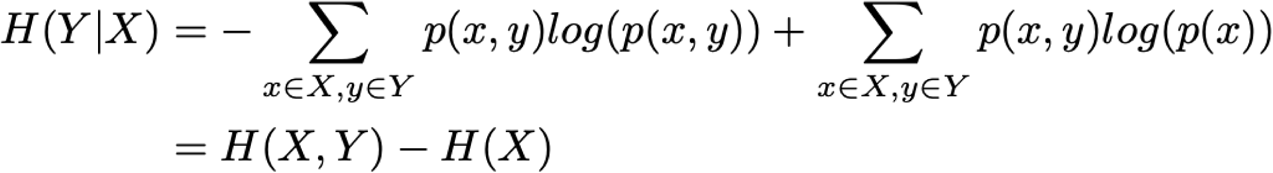

Here p(x) denotes the proportion of times that a particular value of X happens out of all X occurred values in the data set. Similarly, p(x,y) denotes the joint probability of the particular values x, y happening together. For processing the data, we used the Python programming language (version 3.9) (*51*) via Jupyter Notebook (*52*), a web-based interactive computing platform.

Visual attention orienting behavior was coded in a separate pass from maternal behaviors. We calculated here the number of shifts in looking to/from the mother, defined as the number of times annotators indicated that infants clearly returned their gaze to the mother after having looked away. We corrected this value for total time on task for each dyad.

### Glitter Wand Inhibitory Control Task

The child is seated in a caregiver’s lab in front of a small table. The Experimenter shows the child the glitter wand, flash the lights, and move it around. The Expermenter attracts the child’s attention by saying, “look at this wand! Look at its lights!”. They then place the glitter wand on the table within the child’s reach, looks at the child and says “Don’t touch this toy until we say it is Okay. You can play with the toy if you can wait.” The Experimenter looks away from the child and starts a timer once hands have been released from the toy. If 30 seconds passes and the child has not yet touched it, the Experimenter says, “Great job! You didn’t touch it, so now you can play with the toy for a bit!” If the child touches the glitter wand, the timer is stopped and the Experimenter says, “It’s okay; you can touch it now.” The score is the wait time from the release of the glitter wand to the time the child touches it.

### Biospecimen and sequencing

#### Sample Collection

Stool samples were collected at the clinic by a research assistant, who directly transferred them from the diaper to the Zymo DNA/RNA Shield™ Fecal Collection Tube (#R1101, Zymo Research Corp., Irvine, USA) and immediately froze them at -80°C. Samples were not collected if the subjects had taken antibiotics within the preceding two weeks.

#### DNA Extraction

The DNA extraction process was conducted at the Medical Microbiology Department of the University of Cape Town, South Africa. The samples were processed using the Zymo Research Fecal DNA MiniPrep kit (#D4300, Zymo Research Corp., Irvine, USA) according to the manufacturer’s instructions. ZymoBIOMICS® Microbial Community Standards (#D6300 and #D6310, Zymo Research Corp., Irvine, USA) were used as controls and extracted in the same manner as the stool samples. The DNA yield and purity were measured with a NanoDrop® ND2000 spectrophotometer (Thermo Scientific, Nanodrop Technologies Inc., Wilmington, USA).

#### Sequencing

All samples underwent shotgun metagenomic sequencing at the Integrated Microbiome Research Resource (IMR) at Dalhousie University, Nova Scotia, Canada. A pooled library, accommodating up to 96 samples per run, was prepared using the Illumina Nextera Flex Kit for MiSeq and NextSeq from 1 ng of each sample. The samples were then pooled onto a plate and sequenced on the Illumina NextSeq 2000 platform with 150+150 bp paired-end P3 cells, generating 24 million raw reads and 3.6 Gb of sequence per sample (*53*).

#### Metagenome processing

Raw metagenomic sequence reads were processed using tools from the bioBakery as previously described (*54*, *55*). Briefly, KneadData v0.10.0 was used with default parameters to trim lowquality reads and remove human sequences (using reference database hg37). Next, MetaPhlAn v3.1.0 (using database mpa_v31_CHOCOPhlAn_201901) was used with default parameters to map microbial marker genes to generate taxonomic profiles. Taxonomic profiles and raw reads were passed to HUMAnN v3.7 to generate stratified functional profiles.

#### PCI-Microbiome Dataset composition

A paired dataset for analysis of microbial features and Parent-Child Interaction (PCI) outcomes was built from participants who had both metagenomes and recorded/processed videos. Participants were included if the stool sample was collected less than 3 months apart from the video recording. The initial dataset for 94 participants contained the age in days at the time of stool collection, child sex, maternal unpredictability, visual orienting behaviour and the microbial community profiles - both taxonomic and functional - represented as matrices of feature abundances by samples (rows)

#### Stool sample exclusion criterion

One sample was excluded *post hoc* after analysis initially found a significant association between *Streptococcus mitis* abundance and infant VOB. Further inspection revealed that this association was driven by a single outlier with both a high VOB and an abnormally high *S. mitis* relative abundance (>50%). *S. mitis* is typically an oral and respiratory commensal rather than a gut resident, where its presence is linked to dysbiosis and potential pathogenicity (*56*). This sample exerted excessive leverage on the linear model (Cook’s *D* >> 3 × *mean*(*D*)), artificially inflating the effect. Excluding it rendered the *S. mitis*-VOB association non-significant while leaving all other results unchanged. For transparency, data for this sample is included in the Data Availability and Reproducibility statement. As a consequence, the resulting dataset for analysis consisted of 93 paired metagenomes and PCI outcomes.

#### Exploratory microbial community analysis

Principal coordinates analysis was performed in the Julia programming language v1.12.1 (*57*) using the Microbiome.jl package v0.10.0 (*58*). Bray-Curtis dissimilarity was calculated across all pairs of samples, filtering for species-level classification, with Distances.jl v0.10.12. Classical multidimensional scaling was performed on the dissimilarity matrix using MultivariateStats.jl v0.10.3 and PERMANOVA.jl v0.1.1 (*59*).

#### Linear modeling protocol

Individual taxonomic features were assessed for associations with VEP features using multivariable linear models according to the MaAsLin v3 methodology (*60*) implemented using the GLM.jl package v1.8.3 (*61*). The equation tested was species ∼ VOB + mbiome_age_months + child_sex; where species is the arcsin-normalized relative abundance of each taxon when present in the sample, VOB is the unit-range-normalized numerical value of the VOB feature, mbiome_age_months is the child’s age at the time of stool collection, and child_sex is the assigned sex at birth for the participant. FDR correction was employed via a combination of HypothesisTesting.jl v0.11.6 and MultipleTesting.jl v0.6.0 (*62*).

#### Nonlinear modeling protocol

Random Forests (RFs; (*63*) were selected as the nonlinear prediction engine and processed using the DecisionTree.jl v0.12.4 (*64*) implementation, inside the MLJ.jl v0.19.2 (*65*) framework. Models were trained and benchmarked with a repeated cross-validation approach with different random number generator (RNG) seeds. One hundred repetitions of threefold cross-validation (CV) with 10 different intra-fold RNG states each were used, for a total of 3000 experiments per input set. RF hyperparameters were optimized from a grid; gridpoint details are available alongside the data reproducibility notebooks. After the training procedures, the RMSE for VOB, along with Pearson’s correlation coefficient (R), were benchmarked on the validation sets for the winning combination of hyperparameters.

#### Feature Set Enrichment Analysis protocol

To assess the enrichment or depletion of certain microbial feature sets toward PCI outcomes, we performed FSEA analysis on the dataset. First, the binary presence or absence of each gene was modeled via logistic regression inside GLM.jl, including the PCI outcomes (maternal unpredictability, infant visual attention shifts) and control variables (sex, age), using two different models that differ by the inclusion of an interaction term (Equations Model 1, unique; and Model 2, interactive).

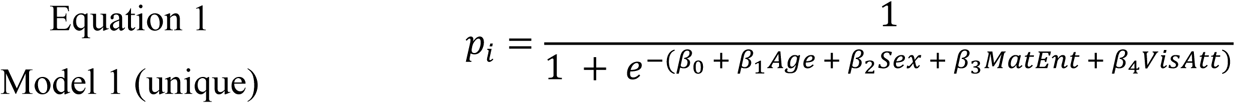

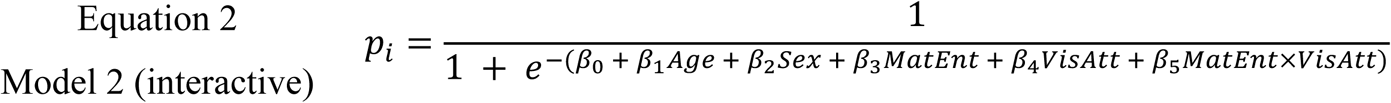

Logistic regression was performed in every individual UniRef90 with all three models. Following that, key individual genes were grouped in gene sets previously curated as mapping to metabolic pathways of neuroactive molecules (*36*). Genes were ranked by the ratio between the parameter estimate and the standard error for each feature of interest. For each neuroactive gene set, enrichment scores (ES) were calculated, and the Wilcoxon rank-sum test was performed to ascertain significant enrichment or depletion (ɑ = 0.05). The multiple-hypothesis correction was performed using the Benjamini-Hochberg procedure, after which gene sets with q < 0.2 were deemed significant (again with HypothesisTesting.jl and MultipleTesting.jl).

## Author contributions

Conceptualization: DA, GFB, VKC

Methodology: DA, GFB, TF, VKC, KAD

Project administration: DA, VKC

Funding acquisition: DA, MG, EM, DCA, DKJ, SCRW, WPF, LJGD, VKC, KAD

Data curation: DA, GFB, TF, KSB, MRZ, FP, MM, DH, CEH, CLD’A, JR

Formal analyses: DA, GFB, MZ

Investigation: DA, GFB

Resources: DA, VKC, KAD

Software and data visualizations: DA, GFB, TF, KSB

Supervision: DA, VKC, KAD

Validation: DA, VKC, KAD

Initial draft: DA, GFB, VKC

Revision and editing: all co-authors

Khula South Africa Data Collection Team contributed to Data curation, Formal analyses and Validation.

## Data and materials availability

All data needed to evaluate the conclusions in the paper are present in the paper and/or the Supplementary Materials. The raw sequencing data for the Khula study have been deposited in the NCBI Sequence Read Archive (SRA) under BioProject accession number PRJNA1128723. Taxonomic and functional microbial profiles, as well as subject demographics necessary for statistical analyses and machine learning, are available on the Data Dryad under DOI 10.5061/dryad.15dv41p9k. Information for replicating the package environment and code for data analysis and figure generation, as well as scripts for automated download of input files, are available on GitHub at https://github.com/Klepac-Ceraj-Lab/PCIVEPMBiome2024 and archived on Zenodo under DOI 10.5281/zenodo.17707109.

## Supplementary Materiasl

**Figure S1.**
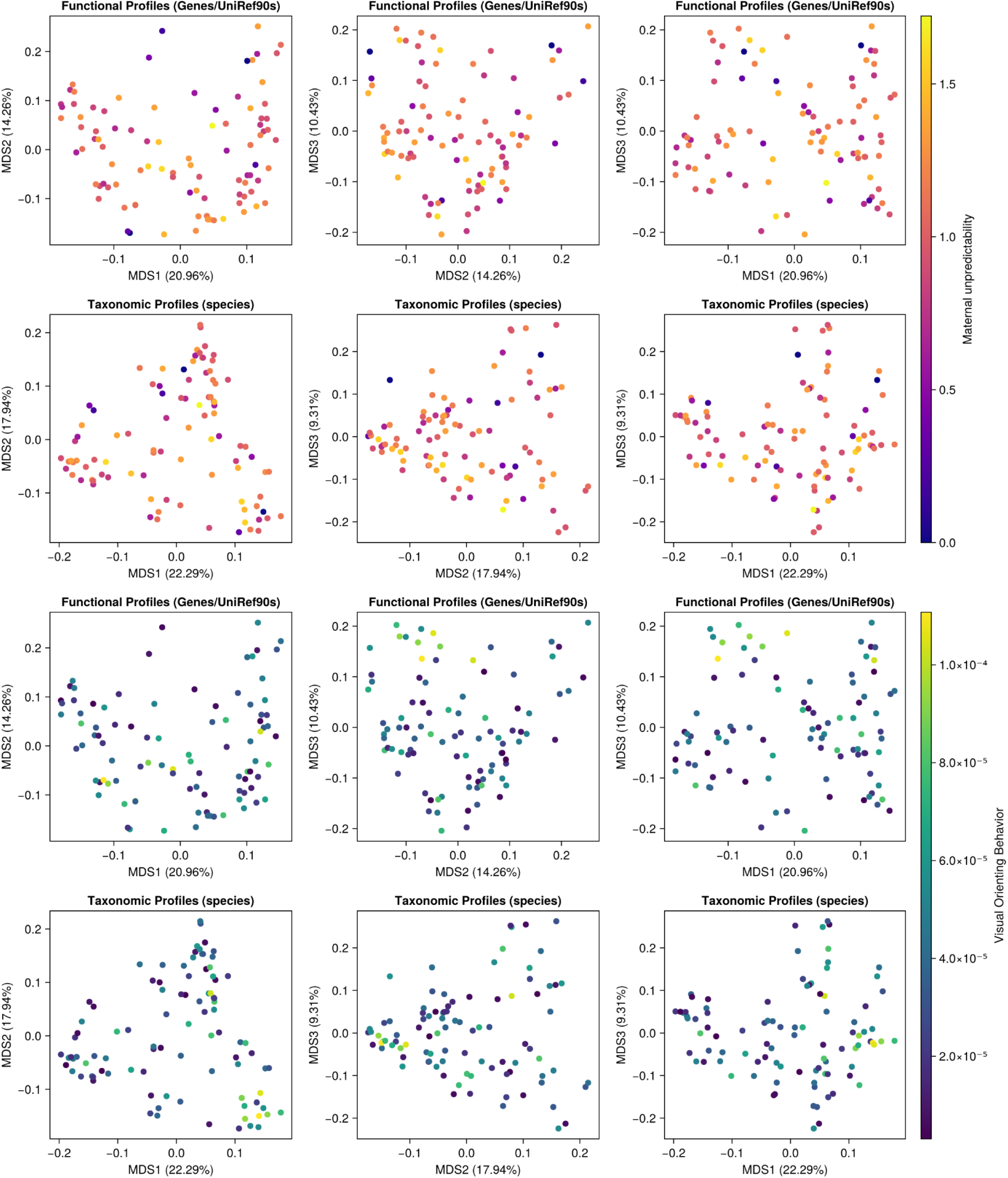
Principal coordinate analysis (PCoA) by non-metric multidimensional scaling (nMDS) on community-wide beta-diversity calculated as Bray-Curtis dissimilarity. Plots display all combinations of PC1, PC2 and PC3, for both taxonomic and functional (UniRef90) profiles. Samples (represented by dots) were colored *a posteriori* by associated PCI interaction metadata: maternal unpredictability (A-F) and infant visual attention shifts (G-L).

**Figure S2.**
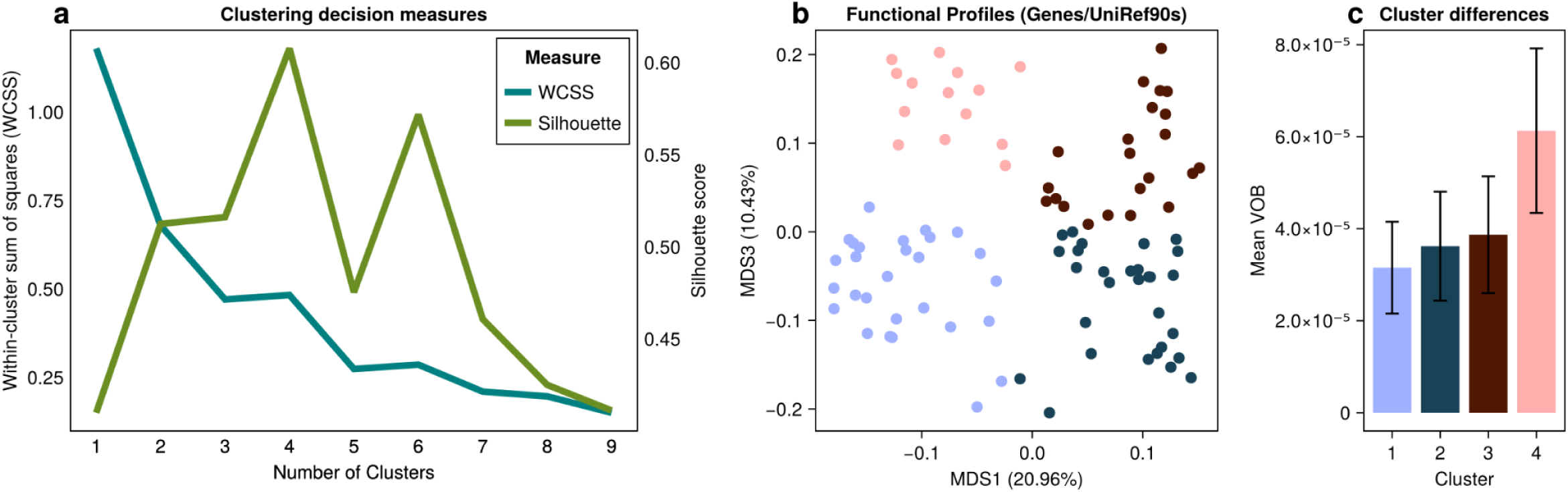
Cluster analysis of microbiome functional profile ordination. (A) Within-Cluster Sum of Squares and Silhouette scores for every k = 1:9. K = 4 was chosen as the best number of clusters due to exhibiting the highest silhouette score with a sufficiently low (<50% of k=1) WCSS. (B) Functional profile PCoA **(Main Figure 3b**) colored by k-means clustering assignment with the selected k=4. (C) Mean and SD of Visual Orienting Behavior in each cluster, illustrating the difference between Cluster 4 (salmon) and clusters 1,2,3.

**Figure S3.**
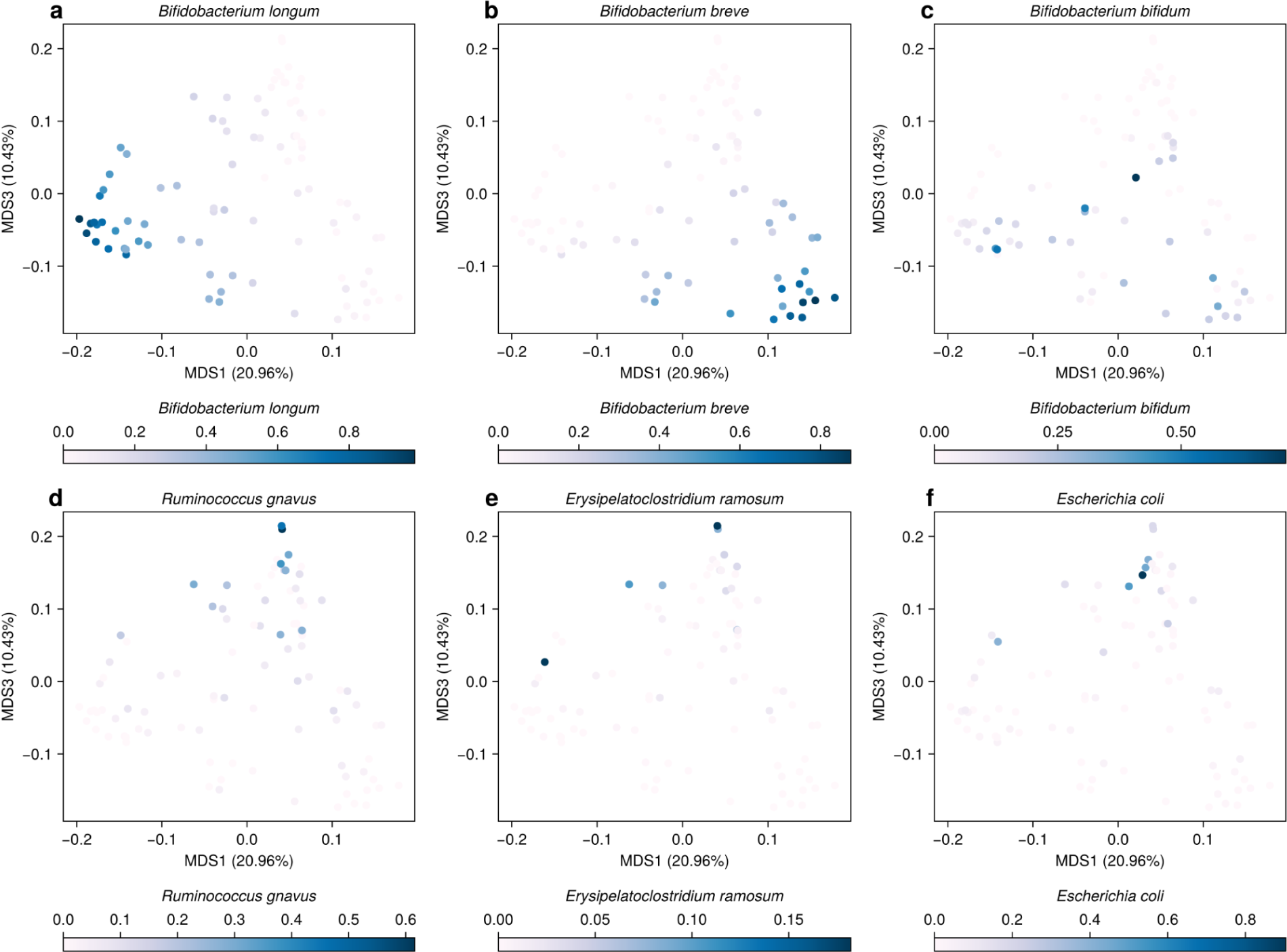
Select taxa distributed along the principal coordinates of the β-diversity ordination of taxonomic profiles. Dots are colored by the relative abundance of a particular taxon. Each taxon is listed above its plot. The percent variance explained is indicated on the X and Y axes. The following taxa are plotted: (A) *Bacterium breve*, (B) *Bacterium longum*, (C) *Bifidobacteirum bifidum*, (D) Ruminococcus gnavus, (E) *Erysipelatoclostridium ramosum*, and (F) *Escherichia coli*.

