## Supplemental Materials for "Microbiome-behavior coupling shapes infant adaptation to early maternal unpredictability"

**This PDF file includes:**

Supplementary Figs. S1 to S3

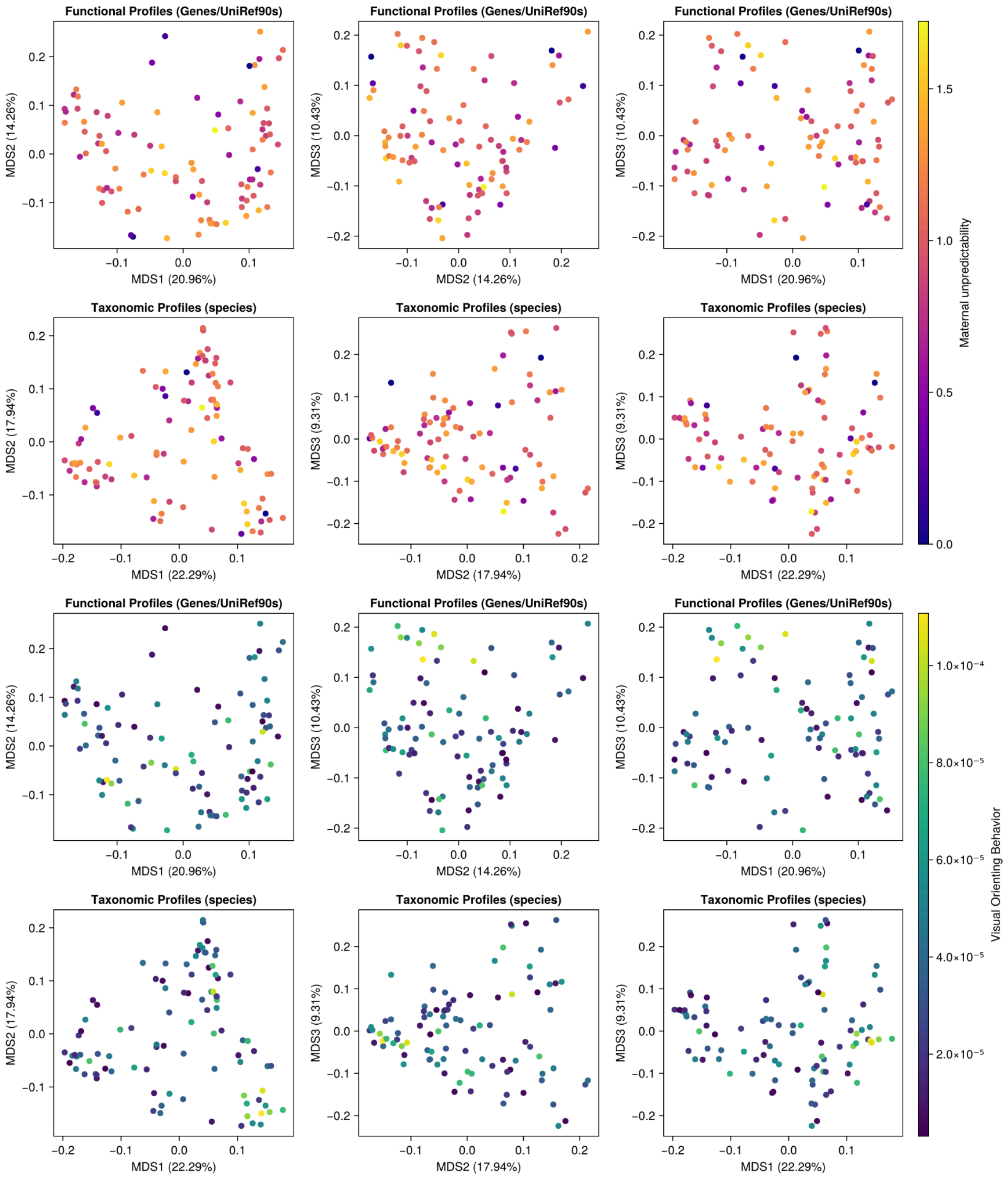


**Figure S1. Principal coordinate analysis (PCoA) by non-metric multidimensional scaling (nMDS) on community-wide beta-diversity calculated as Bray-Curtis dissimilarity.** Plots display all combinations of PC1, PC2 and PC3, for both taxonomic and functional (UniRef90) profiles. Samples (represented by dots) were colored *a posteriori* by associated PCI interaction metadata: maternal unpredictability (A-F) and infant visual attention shifts (G-L).

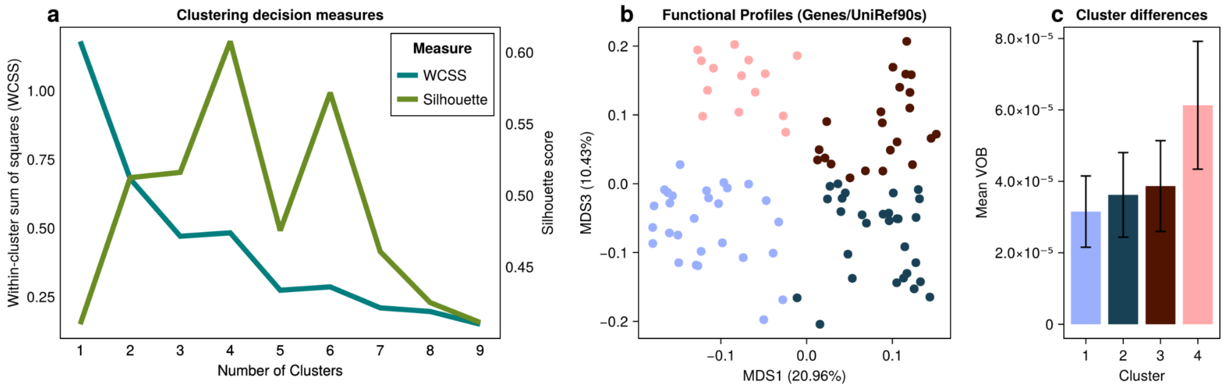


**Figure S2. Cluster analysis of microbiome functional profile ordination.** (A) Within-Cluster Sum of Squares and Silhouette scores for every k = 1:9. K = 4 was chosen as the best number of clusters due to exhibiting the highest silhouette score with a sufficiently low (<50% of k=1) WCSS. (B) Functional profile PCoA **(Main Figure 3b)** colored by k-means clustering assignment with the selected k=4. (C) Mean and SD of Visual Orienting Behavior in each cluster, illustrating the difference between Cluster 4 (salmon) and clusters 1,2,3.

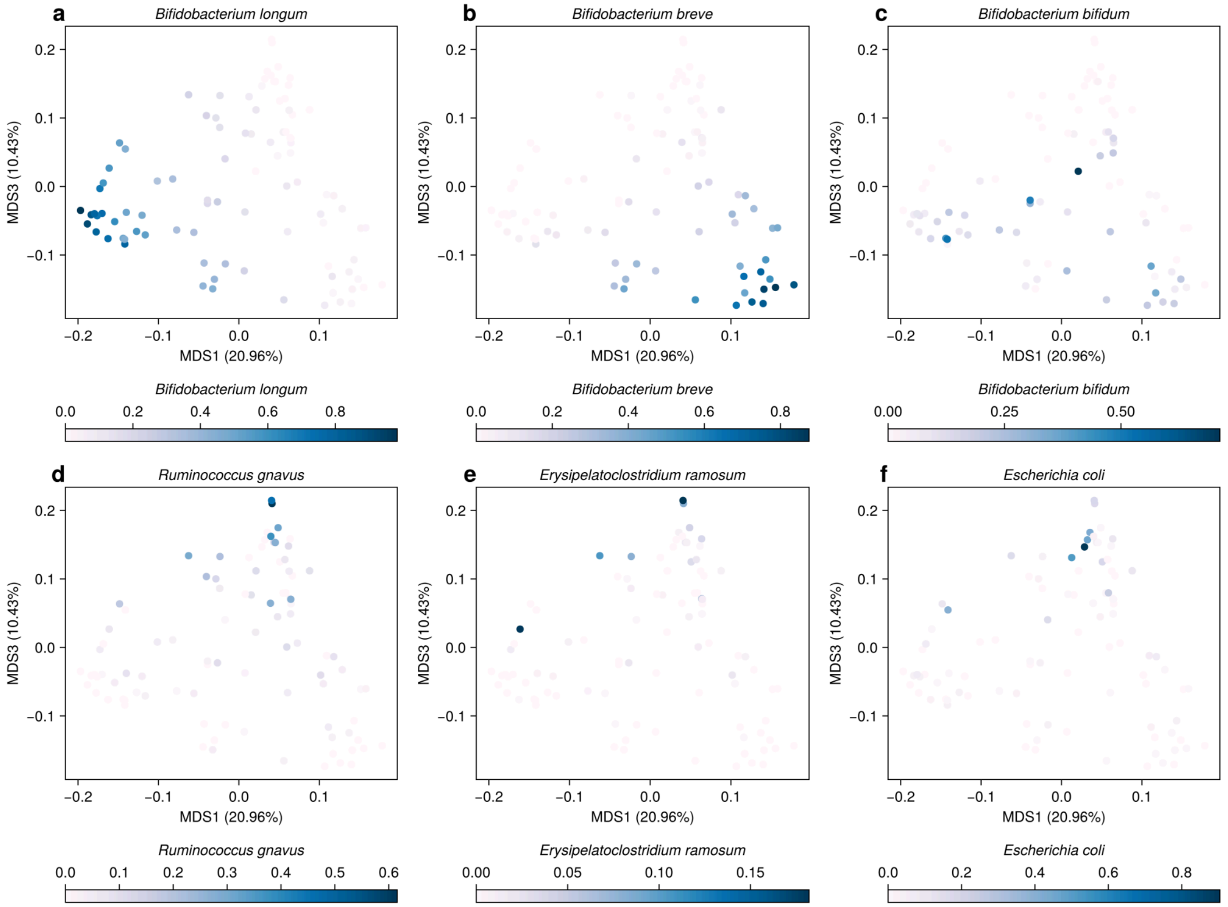


**Figure S3. Select taxa distributed along the principal coordinates of the β-diversity ordination of taxonomic profiles.** Dots are colored by the relative abundance of a particular taxon. Each taxon is listed above its plot. The percent variance explained is indicated on the X and Y axes. The following taxa are plotted: (A) *Bacterium breve*, (B) *Bacterium longum*, (C) *Bifidobacteirum bifidum*, (D) Ruminococcus gnavus, (E) *Erysipelatoclostridium ramosum*, and (F) *Escherichia coli*.
